# Prop3D: A Flexible, Python-based Platform for Machine Learning with Protein Structural Properties and Biophysical Data

**DOI:** 10.1101/2022.12.27.522071

**Authors:** Eli J. Draizen, John Readey, Cameron Mura, Philip E. Bourne

## Abstract

**Background:** Machine learning (ML) has a rich history in structural bioinformatics, and modern approaches, such as deep learning, are revolutionizing our knowledge of the subtle relationships between biomolecular sequence, structure, function, dynamics and evolution. As with any advance that rests upon statistical learning approaches, the recent progress in biomolecular sciences is enabled by the availability of vast volumes of sufficiently-variable data. To be useful, such data must be well-structured, machine-readable, intelligible and manipulable. These and related requirements pose challenges that become especially acute at the computational scales typical in ML. Furthermore, in structural bioinformatics such data generally relate to protein three-dimensional (3D) structures, which are inherently more complex than sequence-based data. A significant and recurring challenge concerns the creation of large, high-quality, openly-accessible datasets that can be used for specific training and benchmarking tasks in ML pipelines for predictive modeling projects, along with reproducible splits for training and testing.

**Results:** Here, we report ‘Prop3D’, a platform that allows for the creation, sharing and extensible reuse of libraries of protein domains, featurized with biophysical and evolutionary properties that can range from detailed, atomically-resolved physicochemical quantities (e.g., electrostatics) to coarser, residue-level features (e.g., phylogenetic conservation). As a community resource, we also supply a ‘Prop3D-20sf’ protein dataset, obtained by applying our approach to CATH. We have developed and deployed the Prop3D framework, both in the cloud and on local HPC resources, to systematically and reproducibly create comprehensive datasets via the Highly Scalable Data Service (HSDS). Our datasets are freely accessible via a public HSDS instance, or they can be used with accompanying Python wrappers for popular ML frameworks.

**Conclusion:** Prop3D and its associated Prop3D-20sf dataset can be of broad utility in at least three ways. Firstly, the Prop3D workflow code can be customized and deployed on various cloud-based compute platforms, with scalability achieved largely by saving the results to distributed HDF5 files via HSDS. Secondly, the linked Prop3D-20sf dataset provides a hand-crafted, already-featurized dataset of protein domains for 20 highly-populated CATH families; importantly, provision of this pre-computed resource can aid the more efficient development (and reproducible deployment) of ML pipelines. Thirdly, Prop3D-20sf’s construction explicitly takes into account (in creating datasets and data-splits) the enigma of ‘data leakage’, stemming from the evolutionary relationships between proteins.

## Introduction

The recent advent of deep learning approaches such as AlphaFold2 [1] now enables one to access the 3D structure of virtually any protein sequence. As was the case for sequence-level data in the 1980s-2000s, 3D structural data on proteins has now been transformed into a readily available commodity. How might such a wealth of structural data inform our understanding of biology’s central *sequence ↔ structure ↔ function* paradigm? Two new, post-AlphaFold2 challenges can be identified: (i) elucidating the *relationships* between all structures in the protein universe, and (ii) armed with millions of new protein structures [2], exploring the limits of protein *function prediction*. Arguably, classic structural bioinformatics paradigms and approaches, which are largely founded on comparative structural analyses, should now be an even more powerful tool in analyzing and accurately predicting protein function.

In structural bioinformatics, the ‘data’ center around biomolecular 3D structures. Here, we take such ‘data’ to mean the geometric structures themselves, augmented (or *featurized*) by a possible multitude of other properties. These other properties can be (i) at potentially varying length-scales (atomic, residue-level, domains, etc.), and (ii) of numerous types, either *biological* in origin (e.g., phylogenetic conservation at a site) or *physicochemical* in nature (e.g., hydrophobicity or partial charge of an atom, concavity of a patch of surface residues, etc.). A significant and persistent challenge in developing and deploying ML workflows in structural bioinformatics concerns the availability of large, high-quality, openly-accessible datasets that can be (easily) used in large-scale analysis and predictive modeling projects. Here, ‘high-quality’ implies that specific training and benchmarking tasks can be performed reproducibly and without undue effort, and that the data-splits for model training/testing/validation are reproducible. A stronger requirement is that the split method also be at least semi-plausible, or not nonsensical, in terms of the underlying biology of a system—e.g., taking into account evolutionary relationships that muddle the assumed (statistical) independence of the splits. (This topic of evolutionary ‘data leakage’, and how we handle it, is presented in detail below.)

A common task in classical bioinformatics involves transferring functional annotations from a well-characterized protein to a protein of interest, if given sufficient shared evolutionary history between the two proteins. A conventional approach to this task typically applies sequence or structure comparison (e.g., via BLAST [3] or TM-Align [4], respectively) of a protein of interest to a database of all known proteins, followed by a somewhat manual process of ‘copying’ or grafting the previously annotated function into a new database record for the protein of interest. However, in the era of ML one can now try to go automatically and more directly from sequence or structure to functional annotations: an ML model can ‘learn’ these evolutionary relationships between proteins as part of the model, thereby obviating the more manual/tedious (and subjective) alignment-related steps.

However, ML workflows for working with proteins—and, in particular, protein 3D structures—are far more challenging, from a technological and data-engineering perspective, than are many of the standard and more routine ML workflows designed to handle inputs in other ML application domains (e.g., for processing images or text). Protein structures are more difficult to work with, from both a basic and applied ML perspective, for several types of reasons, including: (i) fundamentally, all proteins are related at some level through evolution, thereby causing ‘data leakage’ [5]; (ii) raw/unprocessed protein structures are not always biophysically and chemically well-formed (e.g., atoms or entire residues may be missing) [6, 7]; (iii) somewhat related, some protein structures ‘stress-test’ the flexibility and resiliency of existing data structures by having, for instance, multiple rotamers/conformers at some sites; (iv) a protein’s biophysical properties, which are not always included and learned in existing ML models, are just as critical, if not more so, as the raw 3D geometry itself; and (v) there are many different possible representational approaches/models of protein structures (volumetric data, contact-based graphs, etc.) that can yield different results. In short, protein structural data must be carefully inspected and processed before they can be successfully used and split in precise, sensible ways in order to create robust ML models.

Motivated by these challenges, this work presents ‘Prop3D’ and an accompanying resource called ‘Prop3D-20sf’, shown schematically in Figure 1. As a new Python-based platform for processing and otherwise manipulating protein domain structures, Prop3D includes tools to build one’s own datasets with (i) cleaned/prepared structures, (ii) pre-calculated biophysical and evolutionary properties and (iii) different protein representations, alongside (iv) ML-ready train/test splits. We supply the methods by which anyone can readily recreate the Prop3D-20sf dataset supplied with Prop3D; these calculations can be done in a distributed manner and read into the Prop3D framework for use in one’s own ML models. The pre-computed datasets that we provide, using HSDS, can be freely accessed via a standard representational state transfer (ReST) application programming interface (API), along with accompanying Python wrappers for NumPy and the popular ML framework PyTorch. In what follows, we describe the Prop3D software and Prop3D-20sf dataset after first delineating some of the specific considerations that motivated and shaped Prop3D’s design.

**Figure 1:**
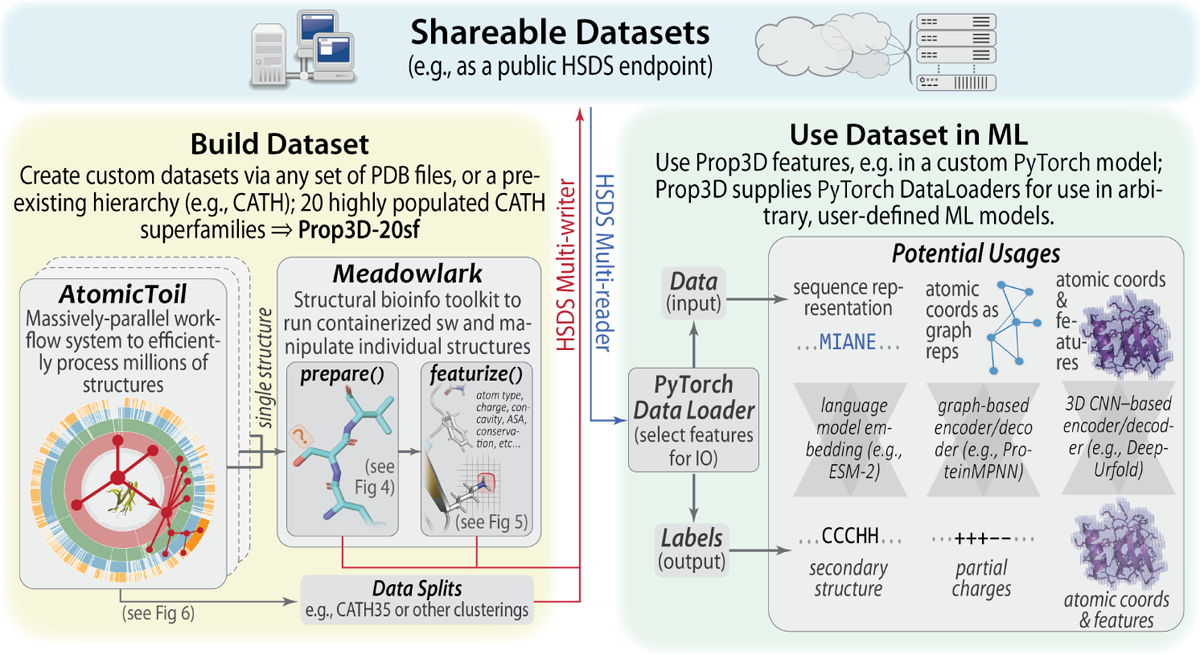
Overview of Prop3D and Its Components. Prop3D is a framework to create and share protein structures featurized with custom sets of properties (biophysical, phylogenetic, etc.), thereby providing ML-ready datasets for structural bioinformatics. One works towards this goal, represented by the green- and blue-background regions to the right and top of this schematic, by utilizing two distinct packages that lie at the core of Prop3D (yellow region at left): (i) ‘Meadowlark’, which enables one to prepare structures, compute and apply features, and run bioinformatics tools/utilities as Docker-ized software (sw); and (ii) ‘AtomicToil’, for performing massively-parallel calculations, locally or in the cloud, using the Toil pipeline system. Proceeding in this way, a dataset of featurized structures can be readily used in the popular ML framework PyTorch, for instance using various representational schemes and types of ML models (language models, graphical models, etc.), as shown in the green region at right; Prop3D facilitates these steps by providing custom PyTorch data loaders that enable rapid, high-volume processing. Prop3D-20sf, a dataset that we created by applying Prop3D to CATH, is available as a publicly-available HSDS endpoint.

### Motivating Factors: Data Leakage, Biophysical Properties, & Protein Representations

#### Evolutionary Data Leakage

ML with proteins is uniquely challenging because all naturally occurring proteins are interrelated via the biological processes of molecular evolution [8]. Therefore, randomly chosen train/test splits are not necessarily meaningful, as there are bound to be crossover relationships between proteins (even if only distantly homologous), ultimately leading to overfitting of the ML model. Moreover, the available datasets are biased—they sample the protein universe in a highly non-uniform (or, rather, *non-representative*) manner (Figure 2), which leads to biased ML models. For example, there are simply more 3D structures available in the Protein Data Bank (PDB [9]) for certain protein superfamilies because, for instance, some of those families were of specific (historical) interest to specific laboratories, certain types of proteins are more intrinsically amenable to crystallization (e.g., lysozyme), some might have been disproportionately more studied and structurally characterized because they are drug targets (e.g., kinases), certain protein families were preferentially selected for during evolution [10], and so on. A possible approach to handle these types of inherent biases would be to create training and validation splits that ensure that no pairs of proteins with *≥* 20% sequence identity occur on the same side of the split [11].

**Figure 2:**
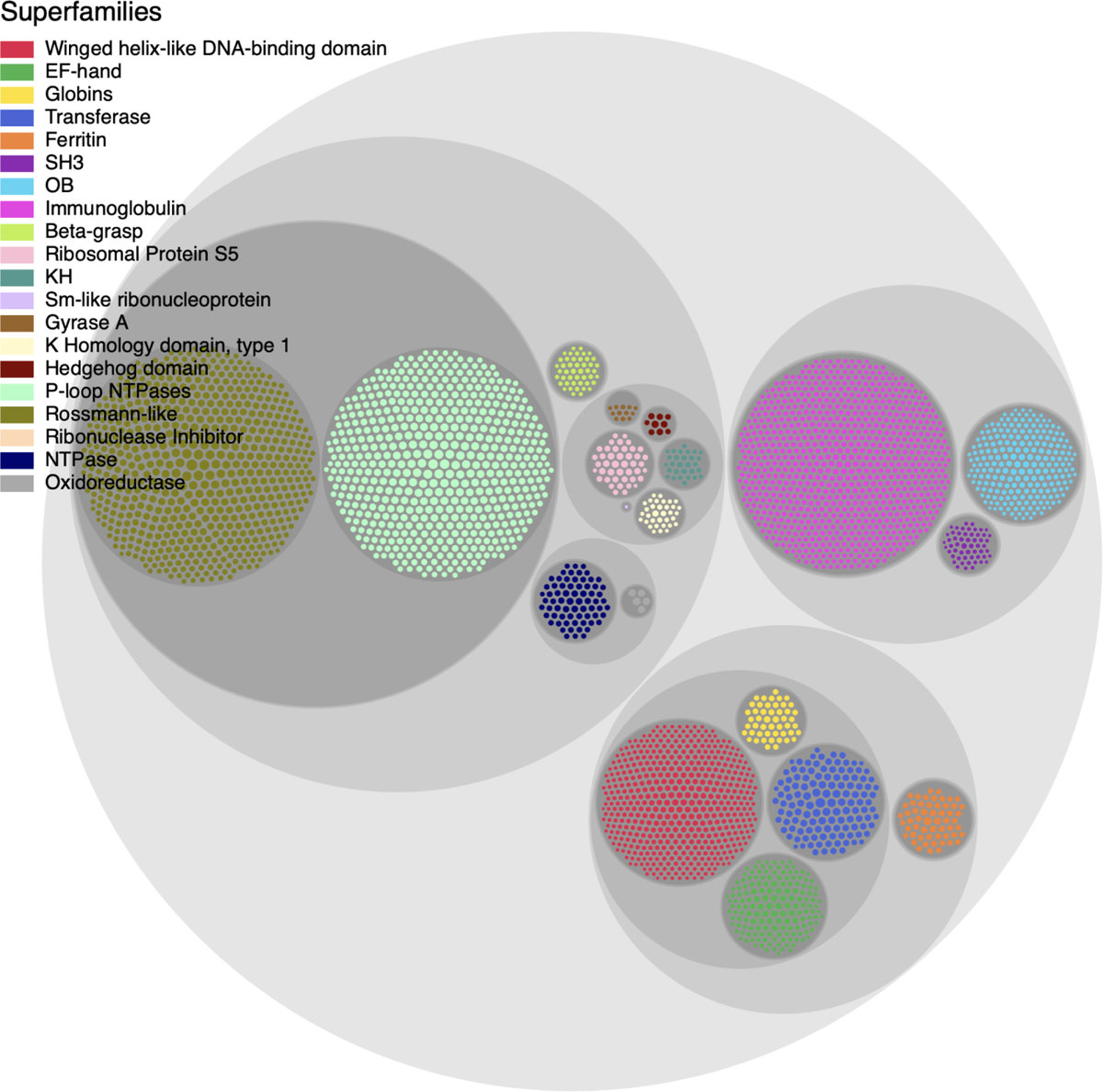
Uneven Distribution of Protein Superfamilies. This diagram of 20 superfamilies of interest, drawn from the CATH hierarchy and shown as a circle-packing diagram, illustrates how the number of known structural domains can vary greatly amongst superfamilies. For instance, superfamilies containing immunoglobulin (magenta), Rossmann-like (olive) and P-loop NTPase (light green) domains are highly abundant versus, e.g., oxidoreductase domains (grey, near center). The Prop3D-20sf dataset is comprised of these 20 highly-populated CATH superfamilies. Possible approaches to mitigate these types of subtle biases would be to (i) create ‘one-class’ superfamily-specific models; or (ii) create multi-superfamily models, making sure to (a) over-sample proteins from under-represented classes and (b) under-sample proteins from over-represented classes [12].

In training ML models at the level of full, intact protein chains, another source of bias in constructing training and validation sets stems from the phenomenon of domain re-use. This is an issue because many full-length protein chains are multidomain (particularly true for polypeptides ≳ 120-150 residues), and many of those individual domains can share similar 3D structures (and functions) and be grouped, themselves, into distinct superfamilies. To illustrate the complexities that must be considered, note that some multi-domain proteins contain multiples of a given protein domain, and the replicates might be virtually identical or highly homologous; in other words, full-length proteins generally evolved so as to utilize individual domains in a highly modular manner (Figure 3). While assigning domains into groups based on an *≈* 20% sequence identity threshold does limit this problem to some extent (if two domains have less than that level of similarity but are still from the same superfamily), a simple, straight-ahead split at 20% identity (or whatever threshold) might negatively impact an ML algorithm at the very basic level of model training. In principle, note that this problem of re-use could also hold at the finer scale of shared structural fragments (i.e., sub-domain–level) too, giving rise to an even more complicated problem.

**Figure 3:**
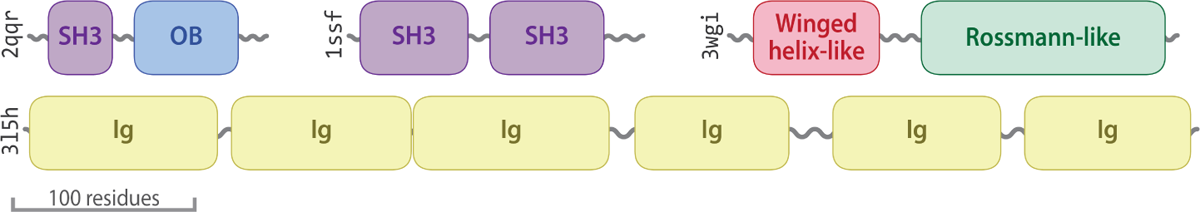
Data Leakage and Multi-domain Proteins. A prime example of evolutionarily-induced data leakage stems from the modular anatomy of many proteins, wherein multiple copies (which often vary only slightly, e.g. as paralogs) of a particular domain are stitched together into a full-length protein. This type of phenomenon is particularly prevalent among protein homologs from more phylogenetically recent species (e.g., eukaryotes like human or yeast, versus archaeal or bacterial lineages). Notably, many proteins that contain SH3, OB and Ig domains are found to include multiple copies of those domains. Examples are schematically illustrated here, using PDB entries 2QQR, 1SSF, 3WGI, and 3L5H.

#### Biophysical Properties in ML

In many ML problems on proteins, it is useful to include biophysical properties mapped onto 3D locations of atoms and residues, thus providing a learning algorithm with additional types of information. However, such properties are often ignored, as in purely sequence-based methods, which neglect 3D structure entirely and frequently use only a one-hot encoding of the sequence, perhaps augmented with some evolutionary information. In other cases, 3D structures are used and only the raw geometry of the atomic structure is used as input, neglecting the crucial biophysical properties that help define a protein’s biochemical properties and physiological functions. There is also a trend in ML wherein one lets a model create its own embeddings, using only a small amount of hand-curated data (e.g., only atom type). Such approaches are generally taken because (i) it is expensive to calculate a full suite of biophysical properties for every atom, say on the scale of the entire PDB (*≈*200K structures); and (ii) the available models, theories and computational formalisms used to describe the biophysical properties of proteins (e.g., approximate electrostatics models, such as the generalized Born) may be insufficiently accurate, thereby adversely influencing the resultant ML models.

Irrespective of the specific details of one use-case or set of tasks versus another, we have found it useful to have available a database of pre-calculated biophysical properties. Among other benefits, such a database would: (i) save time during development of the ML training process, by avoiding repetition of calculations that many others in the community may have already performed on exactly the same proteins (note that this also speaks to the key issue of reproducibility of an ML workflow or bioinformatics pipeline); and (ii) enable one to compare the predicted embeddings of the ML model to known biophysical properties, thereby providing a way to assess the accuracy and veracity of the ML model under development, as well as guide its refinement. Some existing protein feature databases offer various biophysical properties of proteins at different structural ‘levels’ (atomic, residue-based, etc.), as shown in Table 1.

**Table 1:**
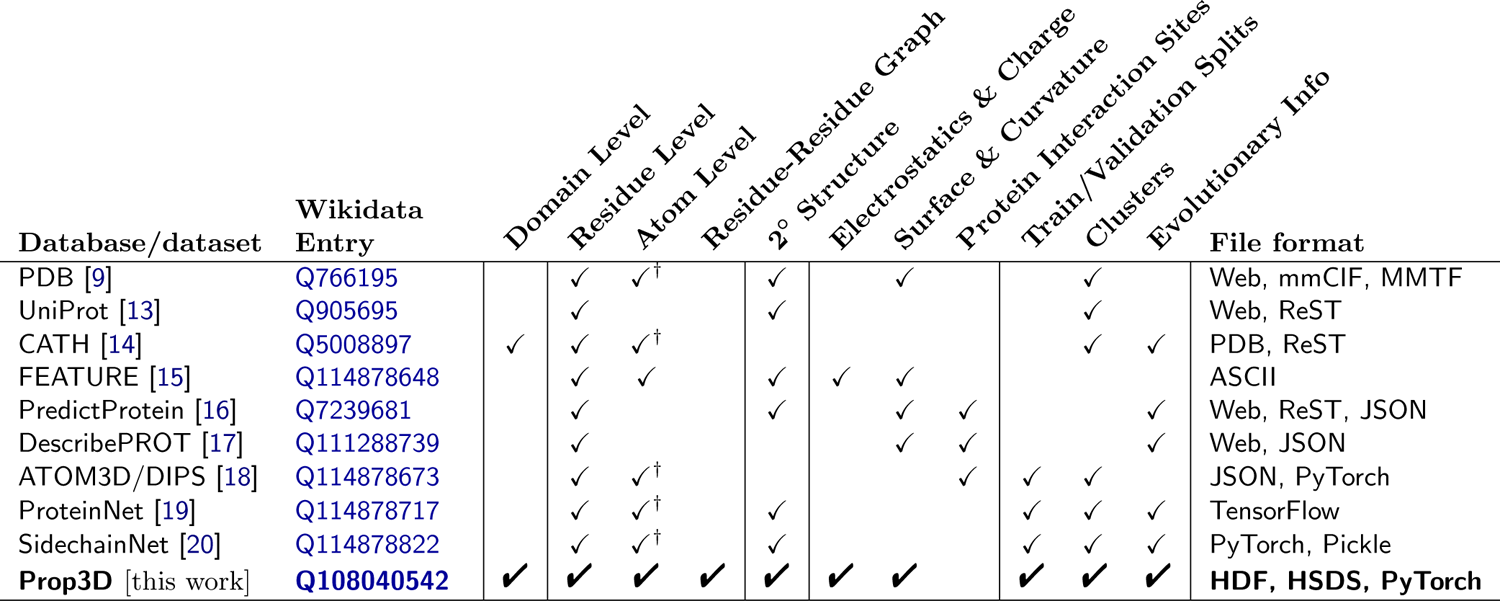
Sequence-based bioinformatics tools available in Prop3D. . Most of these tools have been dockerized, and are available at our Docker Hub (https://hub.docker.com/u/edraizen).

#### Protein Representations

There are various ways to computationally represent a protein for use in ML, each with relative strengths and weaknesses. Many protein structure & feature databases are ‘hard-wired’ so as to include data that can populate only one type of representation; however, to be flexible and agile (and therefore more usable), new databases and database-construction approaches need to allow facile methods to switch between various alternate representations of proteins—i.e., we seek *extensible* structural representation schema. The remainder of this section describes approaches that have been used (Table 2), wherein a protein is represented as a simple sequence, as a graph-based model (residue···residue contact networks), or as a 3D volumetric dataset. We now briefly consider each of these in turn.

**Table 2:**
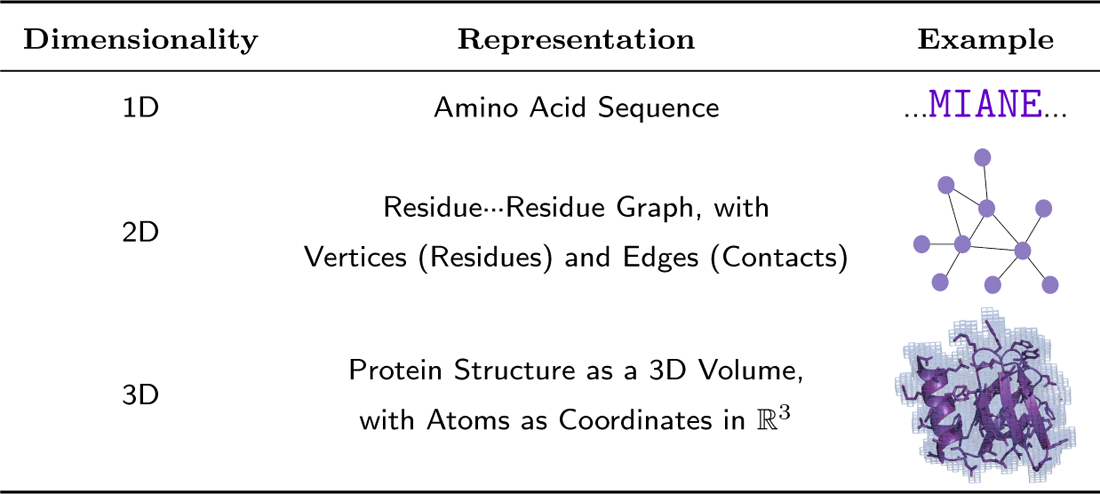
Structural bioinformatics software suites available in Prop3D. . Most of these tools have been dockerized, and are available at our Docker Hub (https://hub.docker.com/u/edraizen).

| Dimensionality | Representation | Example |
| --- | --- | --- |
| 1D | Amino Acid Sequence | ...MIANE... |
| 2D | Residue...Residue Graph, with Vertices (Residues) and Edges (Contacts) |  |
| 3D | Protein Structure as a 3D Volume, with Atoms as Coordinates in $\mathbb{R}^3$ | |

### Proteins in 1D: Sequences

The pragmatically simplest approach to represent a protein is to treat it as a sequence of amino acids, ignoring all structural information (Table 2). In ML workflows, the sequence is generally ‘one-hot encoded’, meaning that each individual character(/residue) in the string is attributed with a 20-element vector; in that vector, all elements are set to zero save the index of the amino acid type that matches the current position, which is set to one. Biophysical properties can also be appended to such representations, giving a feature vector.

### Proteins in 2D: Residue···Residue Graphs

A conceptually straightforward way to capture a protein 3D structure is to build a graph (Table 2), treating the amino acid residues as vertices and interatomic contacts between those residues (near in 3D space) as edges. Individual nodes can be attributed with the one-hot encoded residue type along with biophysical properties, and to each edge can be attributed geometric properties such as a simple Euclidean distance (e.g., between the two residues/nodes), any arbitrary angle of interest (defined by three atoms), any dihedral angles that one likes (defined by four atoms), and so on. These graphs can be fully connected, i.e., with all residues connected to one another, or they may include edges only between residues that lie within a certain cutoff distance of one another (e.g., a 5°A limit to capture van der Waals contacts and other noncovalent interactions).

### Proteins in 3D: Structures as 3D Volumetric Data

Another approach to handle a protein structure in ML is to treat it as a spatially discretized 3D image, wherein volumetric elements (*voxels*) that intersect with an atom are attributed with biophysical properties of the overlapping atom. Here, note that one must define ‘*an atom*’ precisely—e.g. as a sphere of a given van der Waals radius, centered at a specific point in space (the atom’s coordinates), such that the notion of “*intersection* with a specific voxel” is well-defined. Early work in deep neural nets used these types of structural representations, though volumetric approaches have been less prevalent recently for reasons that include: (i) size constraints, with large proteins consuming much memory (scaling with the cube of protein size, in terms of number of residues); (ii) mathematical considerations, such as this representation’s lack of rotational invariance (e.g., structures are manually rotated); (iii) fixed-grid volumetric models are inherently less flexible than graph representations (e.g., 3D images are static and cannot easily incorporate fluctuations, imparting a ‘brittleness’ to these types of data structures); and (iv) related to the issue of brittleness, there exists a rich and versatile family of graph-based algorithms, versus more limited (and less easily implemented) approaches for discretized, volumetric data.

Nevertheless, 3D volumetric approaches, such as are included in Prop3D, have at least two benefits: (i) As long as the complexity is managed [21], 3D representations offer a quite natural way for humans to visualize a protein structure and ‘hold’ the object in mind for analysis [22], versus even 2D graph-based approaches. (ii) The form taken by the data in a 3D volumetric representation is more amenable to explainable AI/ML approaches, such as layer-wise relevance propagation [23], whereby any voxels identified by the algorithm as being ‘important’ can be readily mapped back to specific atoms, residues, patches, etc. in the 3D structure (and those regions may, in turn, be of biochemical or functional interest); such operations are not as readily formulated with 1D (sequence) or 2D (graph) representation schemes. A common approach to voxelize a protein structure into a dense grid is to calculate the distance of every atom to every voxel, then use a Lennard-Jones potential to map scaled biophysical properties to each voxel [24, 25]. This method is feasible for small proteins, but can take an excessively long time for larger structures because of the *O*(*n*^2^) run-time scaling. A faster voxelization approach would be to create a sparse 3D grid, preserving only those voxels that overlap with a van der Waals envelope around each atom; this calculation can be performed using *k* -d trees, with the resultant advantage of scaling as *O*(*n* log *n*) [26, 12].

Finally, note that when treating proteins as 3D images for purposes of training ML models one must take into account the importance of rotational invariance. After translation to a common origin, all protein 3D structures must be repeatedly rotated to achieve (ideally) random sampling of a uniform angular distribution; this task can be viewed in terms of the 3D rotation group *SO*(3), formulated as a Haar distribution over unit quaternions [27]. These numerically-intensive steps add significant computational overhead, thus motivating the pursuit of models that are intrinsically rotationally invariant, e.g., equivariant neural networks [28]. While the data representations for such approaches are not yet pre-built into Prop3D, this is a future direction to consider.

### Birds-eye View of Prop3D and Prop3D-20sf

The remainder of this work presents Prop3D and Prop3D-20sf, the latter of which is a new protein domain structure dataset that includes (i) corrected/sanitized protein 3D structures, (ii) annotated/featurized biophysical properties for each atom and residue, to allow for multiple representation modes, as well as (iii) pre-constructed train, test & validation splits that have been specifically formulated for use in ML of proteins (to mitigate evolutionary data leakage). The tools provided in the Prop3D platform were used to create Prop3D-20sf, for distribution as a community resource.

### Overview of the Software & Associated Dataset

#### Architecture and Design

The Prop3D-20sf dataset is created by using Prop3D in tandem with two other frameworks that we developed: (i) ‘Meadowlark’, for processing and interrogating individual protein structures and (ii) ‘AtomicToil’, for creation of massively parallel workflows of many thousands of structures. An overview of these tools and their relationship to one another is given in Figure 1. While each of these codebases are intricately woven together (in practice), giving the Prop3D functionality, it helps to consider them separately when examining their utility/capabilities and their respective roles in an overall Prop3D-based ML pipeline.

#### Meadowlark: An Extensible, Dockerized Toolkit for Reproducible, Cross-platform Structural Bioinformatics Workflows

In bioinformatics and computational biology more broadly, many tools and codes can be less than straightforward to install and operate locally: They each require particular combinations of operating system configurations, specific versions of different languages and libraries (which may or may not be cross-compatible), have various dependencies for installation/compilation (and for run-time execution), potentially difficult patterns of interdependencies, and so on. Moreover, considered across the community as a whole, researchers spend many hours installing (and perhaps even performance-tuning) these tools themselves, only to find that they are conducting similar development and upkeep of this computational infrastructure as are numerous other individuals. All the while, the data, results and technical/methodological details underpinning the execution of a computational pipeline are typically never shared, at least not before the point of eventual publication— i.e., months or even years after the point at which it would have been most useful to others. Following the examples of the UC Santa Cruz Computational Genomics Laboratory (UCSC-CGL) and the Global Alliance for Genomics & Health (GA4GH) [29], in Prop3D we Docker-ize common structural bioinformatics tools to make them easily deployable and executable on any machine, along with parsers to handle their outputs, all without leaving a top-level Python-based workflow. New software can be added into meadowlark if it exists as a Docker or Singularity container [30, 31]; indeed, much of Prop3D’s extensibility stems from meadowlark, and new functionality can be readily added beyond the provided *prepare()* and *featurize()* tools shown in Figure 1. For a list of codes and software tools that we have thus far made available, see Supplemental Table 1 & Supplemental Table 2 or visit our Docker Hub for the most current information.

#### AtomicToil: Reproducible Workflows That Map Structural Information to Sets of Massively Parallel Tasks

To enable the construction and automated deployment of massively parallel workflows in the cloud, we use a Python-based workflow management system (WMS) known as Toil [30]. Each top-level Toil job has child jobs and follow-on jobs, enabling the construction of complex *MapReduce*-like pipelines. A Toil workflow can be controlled locally, on the cloud (e.g., AWS, Kubernetes), or on a compute farm or a high-performance computing platform such as a Linux-based cluster (equipped with a scheduler such as SLURM, Oracle Grid Engine, or the like). Further information on the data-flow paradigm, flow-based programming and related WMS concepts, as they pertain to task-oriented bioinformatics toolkits such as Toil, can be found in [32].

In Prop3D, we have specifically created multiple ways by which a user can develop and instantiate a workflow. Namely, pipelines can be devised based on:

1. <u>PDB files</u>: A collection of PDB files, each of which can contain a single protein domain or perhaps be more complex (e.g., multiple chains), can be aggregated into a pool. This group of PDB identifiers can be systemically mapped to jobs in order to run a given function/calculation (‘*apply*’ the function, in the parlance of functional programming) on each member of the data pool, thereby processing the full dataset.
2. <u>CATH’s schema</u>: The CATH database is readily amenable to the data-flow paradigm by virtue of its hierarchical organization. In this scheme, one job/task can be created for each *n*^th^ level entry in the CATH hierarchy, with child jobs spawned for subsidiary *n*+1^th^ levels in the hierarchy. Once the workflow reaches a job at the level of each individual domain (or whatever pre-specified target level), then it can run a given, user-provisioned function.

New, user-defined functionality can be added to a workflow by defining new Toil job functions; these functions can be arbitrarily complex, or as simple as standalone Python functions with specific, well-formed signatures (call/return semantics).

#### Capabilities and Features

This section offers two examples of Prop3D usage, one relatively simple and the other more intermediate-level. The more advanced example demonstrates protein structure preparation and biophysical property calculations (and annotation). While not included here, we note that Prop3D is also useful in creating more intricate workflows, for instance (i) to build and validate intermolecular associations, e.g., in studying domain···domain interactions and protein complexes, and (ii) in developing and deploying an AI-driven ‘DeepUrfold’ framework for quantifying protein structural relationships [12].

### Example 1: Protein Structure Preparation

To illustrate the typical first step in a structural bioinformatics analysis pipeline, we ‘clean’ or ‘sanitize’ a starting protein 3D structure via the following scheme. We begin by selecting the first model (from among multiple possible models in a PDB file), the desired chain, and the first alternate location (if multiple conformers exist for an atom/residue). These two choices are justifiable, in the absence of other information, because in the PDB file-format it is conventional for (i) the first ‘MODEL’ to be the lowest-energy (most energetically favorable) conformation, e.g., in NMR-derived structural ensembles or theoretical predictions, and (ii) similarly, the first rotameric state, specified by alternate location (‘altloc’) identifiers, corresponds to the most highly-populated (and presumably lowest-energy) side-chain conformer. Next, we remove hetero-atoms (water or buffer molecules, other crystallization reagents, etc.); these steps are achieved in Prop3D via pdb-tools [33]. Then, in the final phase, we modify each domain structure via the following stages: (i) Build/model any missing residues with MODELLER [34]; (ii) Correct/optimize rotamers (e.g., any missing atoms) with SCWRL4 [35]; and (3) Add hydrogens and perform a rough potential energy minimization with the PDB2PQR toolkit [36]. Again, we note that all these software packages and utilities are wrapped into Prop3D’s unified framework. We applied this general workflow, schematized in Figure 4, in constructing the Prop3D-20sf dataset.

**Figure 4:**
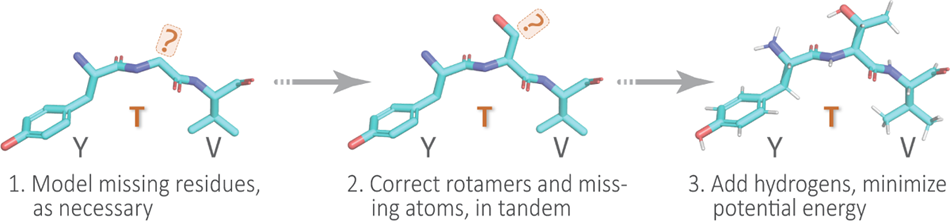
A Simple Protein Preparation Pipeline. In working with protein structures, e.g., to create the Prop3D-20sf dataset, each domain is typically corrected or ‘sanitized’ by adding missing atoms and residues, checking rotameric states (highly-populated rotamers should be assigned, by default), protonating, and performing a crude potential energy minimization of the 3D structure; this general workflow is sketched here using a tripeptide segment (PDB entry 1KQ2).

### Example 2: Biophysical Property Calculation & Featurization

The Prop3D toolkit enables one to rapidly and efficiently compute biophysical properties for all structural entities (atoms, residues, etc.) in a dataset of 3D structures (e.g., from the PDB or CATH), and then map those values onto the respective entities as features for ML model training or other downstream analyses.

For atom-level features, we create one-hot encodings based on 23 atom names, 16 element names, and 21 residue types (20 standard amino acids and one UNKnown placeholder), as defined in AutoDock. We also include van der Waals radii, charges from PDB2PQR [36], electrostatic potentials computed via APBS [37], concavity values that we calculate via CX [38], various hydrophobicity features of the residue that an atom belongs to (the Kyte-Doolittle [39], Biological [40] and Octanol [41] scales), and two measures of accessible surface area (per-atom, via FreeSASA [42], and per-residue, via DSSP [43]). We also include different types of secondary structure information, namely one-hot encodings based on DSSP’s 3-class (helix, strand, loop) and more finely-grained 7-class secondary structure classifications (the latter also includes an eighth class for ‘unknown’/error types), as well as the backbone torsion angles φ and ψ (along with embedded sine and cosine transformations of each). We also annotate aromaticity, and hydrogen-bond acceptors and donors, based on AutoDock atom-name types. As a gauge of phylogenetic conservation, we include sequence entropy scores from EPPIC [44]. These biophysical, physicochemical, structural, and phylogenetic features are summarized in Figure 5 and are exhaustively enumerated in Table 3. Finally, Prop3D also provides functionality to create discretized values of features via the application of Boolean logic operators to the corresponding continuous-valued quantities of a given descriptor, using simple numerical thresholding (Table 4).

**Figure 5:**
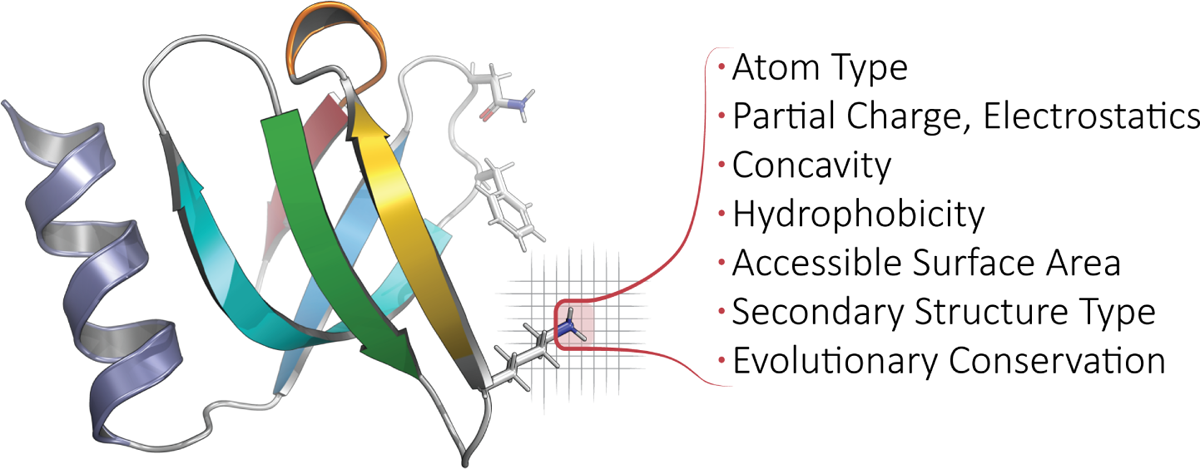
Calculated Properties/Features, Biophysical and Beyond. For each protein domain in Prop3D-20sf, we annotate every atom with the following features: atom type, element type, residue type, partial charge & electrostatics, concavity, hydrophobicity, accessible surface area, secondary structure type, and evolutionary conservation. For a full list of features used in Prop3D-20sf, see the text and Tables 3 & 4. In the ribbon diagram shown here (PDB 1KQ2), a featurized (atomic) region is highlighted and demarcated in red, atop a voxelized background. Note that any bespoke feature can be defined and applied in Prop3D.

**Table 3:**
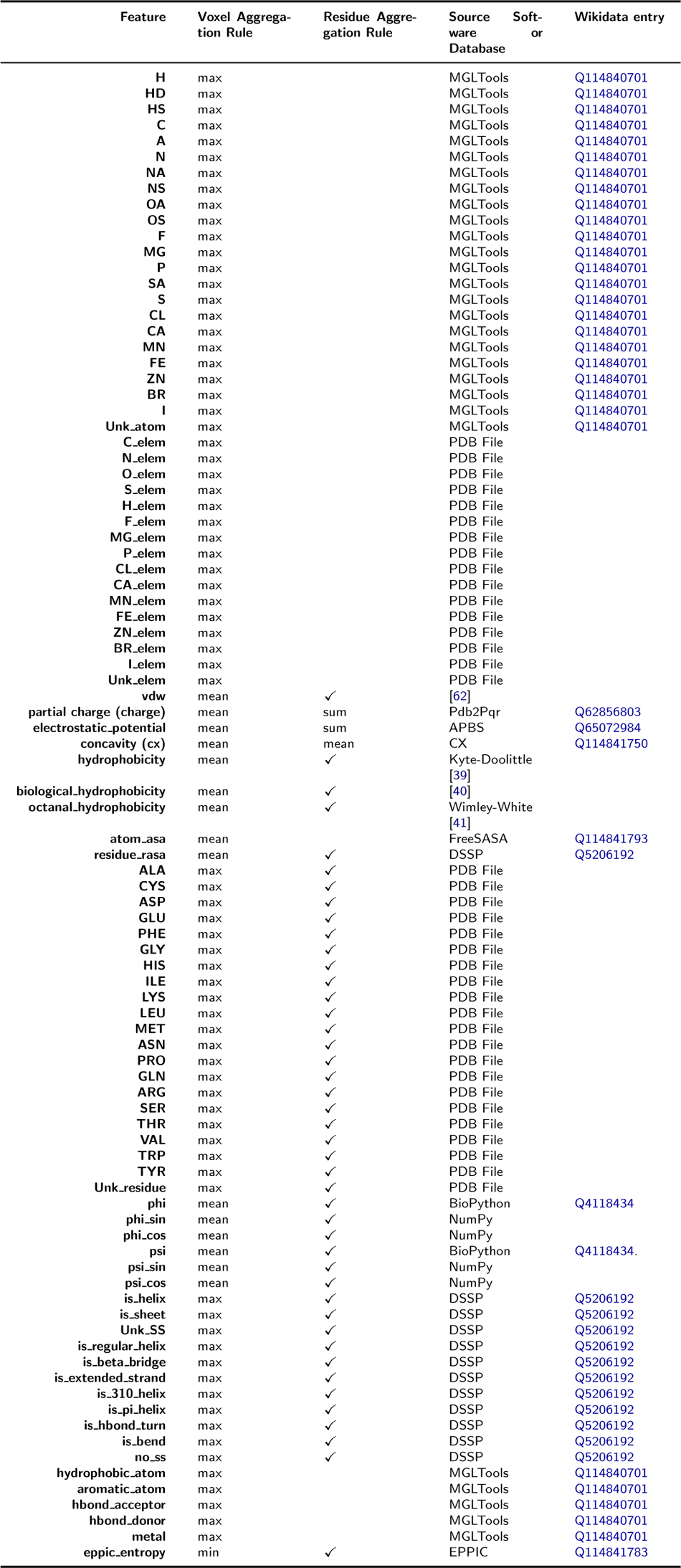
All Protein Descriptors and Extracted Features That Are Defined and Calculated in Prop3D. A voxel aggregation method is used to combine two or more atom-wise features if they impinge upon the same voxel (after accounting for the van der Waals sphere volume). The “Residue Aggregation Rule” describes how a given feature is aggregated, from atom to residue, if also present as a residue-level feature. A ‘✓’ indicates if a feature was calculated at the residue level and mapped down to the atomic level.

**Table 4:**
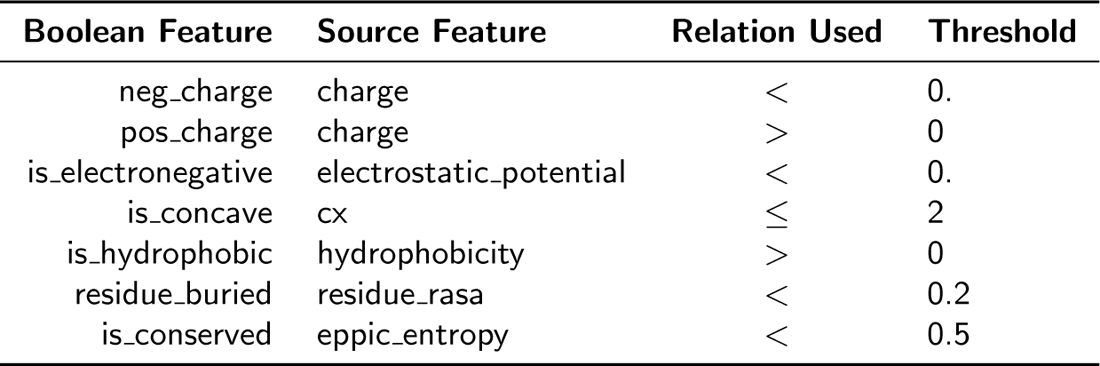
Discretization of Continuous Feature Values in Prop3D. . Boolean operators, and the associated threshold criteria listed here, are applied in order to obtain discretized features by converting from continuous-valued ones.

Some of the properties mentioned above are computed at the residue level and mapped to each atom in the residue (e.g., hydrophobicity is one such property). That is, a ‘child’ atom inherits the value of a given feature from its ‘parent’ residue. For other features, residue-level values are calculated by combining atomic quantities, via various summation or averaging operations applied to the properties’ numerical values (as detailed in Table 3 for Prop3D-20sf). To illustrate the principle that residue-level properties may be directly/simply or indirectly/complexly related to atomic properties, consider that (i) the mass of a residue is a simple summation of the atomic masses of each of its constituent atoms, whereas (ii) properties such as residue volume or accessible surface area are not so straightforwardly derived from atomic properties, instead requiring the application of geometric methods (e.g., the Shrake-Rupley numerical algorithm [45]).

While all of the possible features are contained in the Prop3D-20sf dataset and undoubtedly will be somewhat correlated, it is possible for one to select only certain subsets of features of interest. We also create subsets of the Boolean features that we have found to be minimally correlated [46], and those can be selected, for example, in training deep neural networks.

As illustrative use-cases, we supply three nontrivial ML examples that involve representing proteins as sequences, graphs, or full 3D structures. At the sequence level, we present an example that uses Prop3D together with the language model– based Evolutionary Scale Model approach (ESM-2 [47]) to predict and annotate residue-level properties. Next, we illustrate how Prop3D can be used with Protein-MPNN [48], which is a recent deep learning approach for protein sequence design wherein structural information is encoded as graph neural networks, in order to predict residue-level features. And, finally, we briefly highlight a new DeepUrfold framework [12], where Prop3D is instrumental in creating superfamily-specific deep convolutional variational autoencoder (VAE) models at the level of full, intact 3D structures. These three sets of examples (complete with Python code), along with much other documentation, can be found at https://prop3d.readthedocs.io.

### Dataset Design and Open Data Format (with Some Historical Context)

In order to handle the large amount of protein data in massively parallel workflows, we engineered Prop3D to employ the Hierarchical Data Format (HDF5 [49]), along with the Highly Scalable Data Service (HSDS). We find the HDF5 file format to be a useful way to store and access immense protein datasets because it allows Prop3D to chunk/compress/navigate a protein structure hierarchy like CATH in a scalable and efficient manner. Using this approach versus, for example, creating myriad individual files spread across multiple directories, we can combine the data into ‘single’ files/objects that are easily shareable and can be accessed via a hierarchical structure of groups and datasets, each with attached metadata descriptors; note that hierarchical schemes, such as CATH, will generally lend themselves naturally to this sort of approach. Moreover, the HSDS extension to this object storage system allows multiple readers and writers which, in combination with Toil, affords a degree of parallelization that significantly accelerates the creation of new datasets, e.g. as part of a Prop3D-enabled workflow.

Many computational biologists have begun migrating to approaches such as HDF5 [50, 51, 52] and HSDS [53] in recent years because (i) binary data can be rapidly retrieved/read, (ii) such data are readily manipulable and easily shareable, and (iii) these systems provide well-integrated metadata and other beneficial services, schema and features (thus, e.g., facilitating attribution of data provenance). Before the relatively recent advent of HDF5(/HSDS) and other binary formats, biological data exchange and archival formats for protein 3D structures largely relied on human-readable, plaintext ASCII files (i.e., PDB files). For decades, PDB files have been the *de facto* standard format for sharing, storing and processing protein structure data, such as in structural bioinformatics workflows. Originally developed in 1976 to work with punch cards, the legacy PDB format is an ASCII file with fixed-column width and maximally 80 characters per line [54]. Working with traditional PDB files, a structure could be attributed with only one type of biophysical property, e.g., by substituting the numerical values of the desired property into the *B* -factor column—a highly limited workaround. Because of the inflexibility of the legacy PDB file and its limitations as a data exchange format, the macromolecular Crystallographic Information File (mmCIF) was developed; this file format was designed for better extensibility, flexibility and robustness (e.g., a standardized data dictionary), allowing for a 3D structure to be attributed with a plethora of properties, biophysical and otherwise [55]. Most recently, spurred by the slow nature of reading ASCII files, the Macromolecular Transmission Format (MMTF) has been developed to store protein structures in a compact binary format, based on MessagePack format (version 5) [56, 57]. While the MMTF is almost ideal for ML tasks, it still relies on using individual files in a file system, with no efficient, *distributed* mechanism to read in all files, no way to include metadata higher than residue level, and no ability to combine train/test splits directly into the schema—these were some of our motivating factors in adopting HDF5 and HSDS capabilities in Prop3D.

For Prop3D and Prop3D-20sf, an HDF5 file is built by starting with the CATH database, which provides a hierarchical schema—namely, *Class ⊃ Architecture ⊃ Topology ⊃ Homologous Superfamily* —that is naturally amenable to parallelization and efficient data traversal, as shown in Figure 6. In Prop3D, a superfamily can be accessed by its CATH code as the group key (e.g., ‘2/60/40/10’ for Immunoglobulin). We then split each superfamily into two groups (Figure 6): (i) a ‘domains’ dataset, containing groups for each protein domain inside that superfamily (Figure 6B, top half), and (ii) ‘data splits’ (Figure 6B, bottom half), containing pre-computed train (80%), validation (10%), and test (10%) data splits for use in developing ML models, where each domain in each split is hard-linked to the group for that domain (dashed green arrows in Figure 6). Each domain group contains datasets for different types of features: ‘Atoms’, ‘Residues’ and ‘Edges’. The ‘Atoms’ dataset contains information drawn from the PDB file’s ATOM field, as well as all of the biophysical properties that we calculated for each atom. ‘Residues’ contains biophysical properties of each residue and position (average of all of its daughter atoms), e.g. for use in coarse-grained models. Finally, ‘Edges’ contains properties for each *residue ↔ residue* interaction, thereby enabling the construction and annotation of, e.g., contact maps in graph-based representations/models.

**Figure 6:**
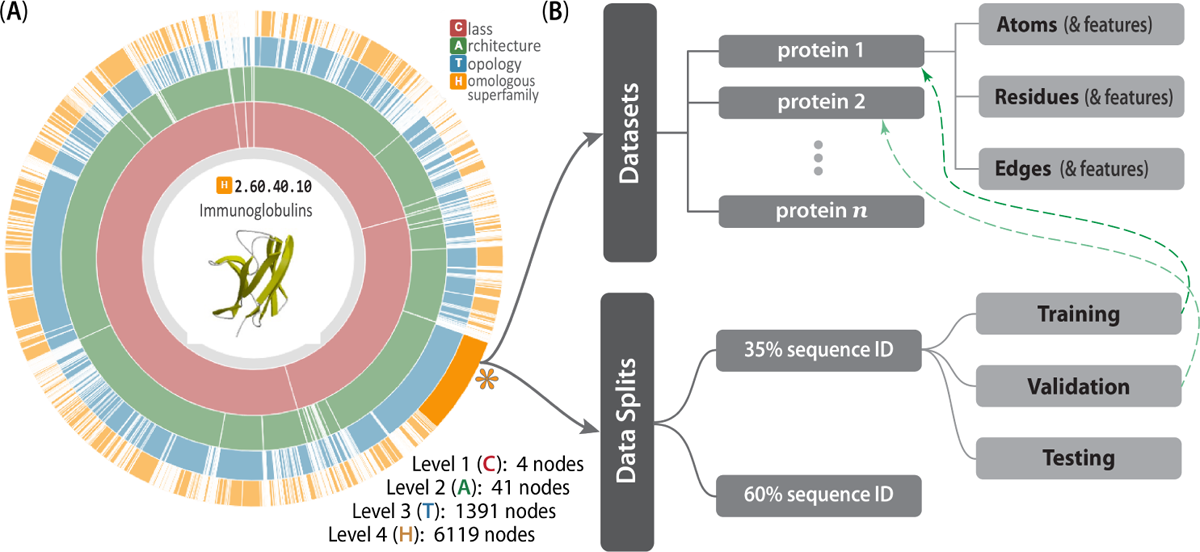
The CATH-inspired Hierarchical Structure of Prop3D. The inherently hierarchical structure of CATH (**A**) is echoed in the design schema underlying the Prop3D-20sf dataset (**B**), as illustrated here. Prop3D can be accessed as an HDF5 file seeded with the CATH hierarchy for all available superfamilies. For clarity, an example of one such superfamily is the individual H-group 2.60.40.10 (Immunoglobulins), shown here as the orange sector (denoted by an asterisk near 4 o’clock). Each such superfamily is further split into (i) the domain groups, with datasets provided for each domain (atomic features, residue features, and edge features), as delineated in the upper-half of (**B**), and (ii) precalculated data splits, shown in the lower-half of (**B**), which exist as hard-links (denoted as dashed green lines) to domain groups. (The ‘sunburst’ style CATH diagram, from http://cathdb.info, is under the Creative Commons Attribution 4.0 International License.)

In terms of data-processing pipelines, HSDS allows HDF5 data stores to be hosted in S3-like buckets, such as AWS or MinIO, remotely and with accessibility achieved via a ReST API. HSDS data nodes and service nodes (Figure 7) are controlled via a load-balancer in Kubernetes in order to enable efficient, distributed mechanisms to query HDF5 data stores, as well as write data with a quick, efficient, distributed mechanism; these properties of HSDS are achieved via various features of its engineering, including using data-caching and implicit parallelization of the task mapping across virtual partitions (Figure 7). HSDS allows for multiple readers and multiple writers to read or write to the same file simultaneously, using a ‘distributed’ HDF5 multi-reader/multi-writer Python library known as h5pyd (Figure 7). As part of Prop3D, we have setup a local k3s instance, which is an easy-to-install, lightweight distribution of Kubernetes that can run on a single machine along with MinIO S3 buckets. We have found this approach to be particularly useful in enabling flexible scalability: our solution works on HPC data infrastructures that can be either large or (relatively) small.

**Figure 7:**
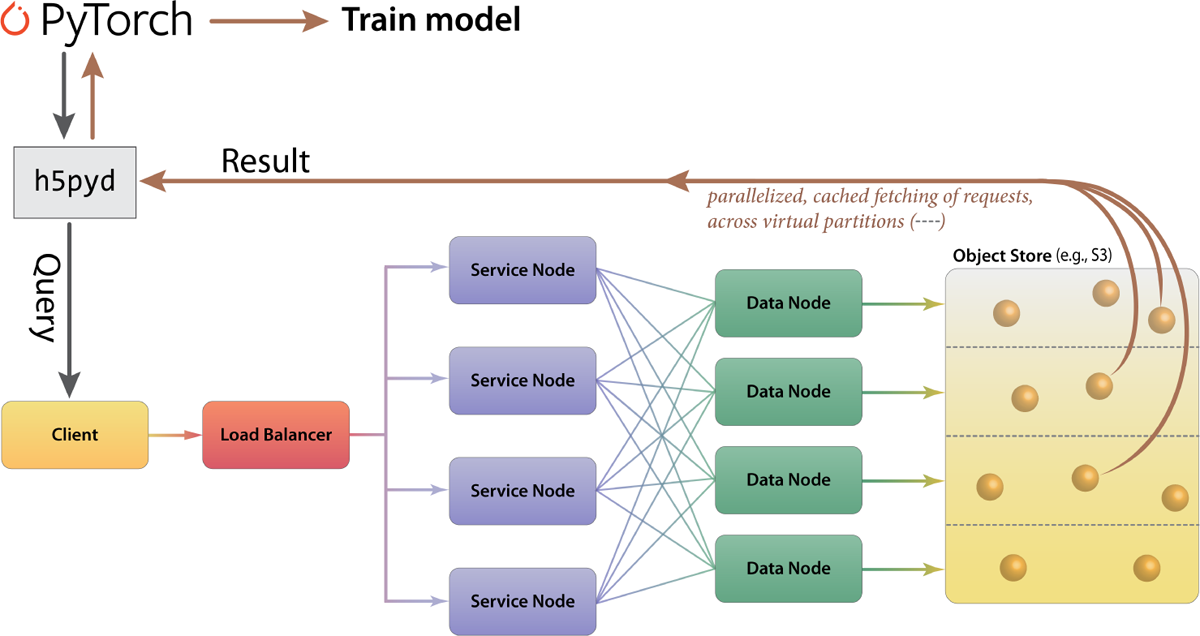
Cloud-based Access to the Prop3D-20sf Dataset via HSDS. HSDS creates Service Nodes, which are containers that handle query requests from clients, and Data Nodes, which are containers that access the object storage in an efficient distributed manner. The Prop3D-20sf dataset can be used as input to train an ML model either by accessing the data via a Python client library (h5pyd) or through our separate DeepUrfold Python package, which supplies PyTorch data loaders [12]. This illustration was adapted from one that can be found at the HSDS webpage (available under an Apache 2.0 license, which is compatible with CC-by-4.0).

In creating the Prop3D-20sf dataset, HSDS, in combination with a Toil-enabled workflow, allows for each parallelized task to write to the same HDF5 data store simultaneously. The Prop3D-20sf dataset can be read in parallel as well, e.g. in PyTorch. We provide PyTorch Data Loaders to read the Prop3D-20sf dataset from an HSDS endpoint using multiple processes; that functionality is available in our related DeepUrfold Python package [12]. Promisingly, we found that when HSDS was used with Prop3D as a system for distributed training of deep generative models in our DeepUrfold ML workflow, as opposed to using raw ASCII files, a speedup of *≈* 33% (8 hours) was achieved, corresponding to a reduction from *≈* 24 hours to *≈* 16 hours of wall-clock time to train an immunoglobulin-specific variational autoencoder model with 25,524 featurized Ig domain structures (Figure 8). Thus, we found it clearly and significantly advantageous to utilize the parallelizable data-handler capacity that is provided by a remote, cloud-based, parallel-processing system like HSDS.

**Figure 8:**
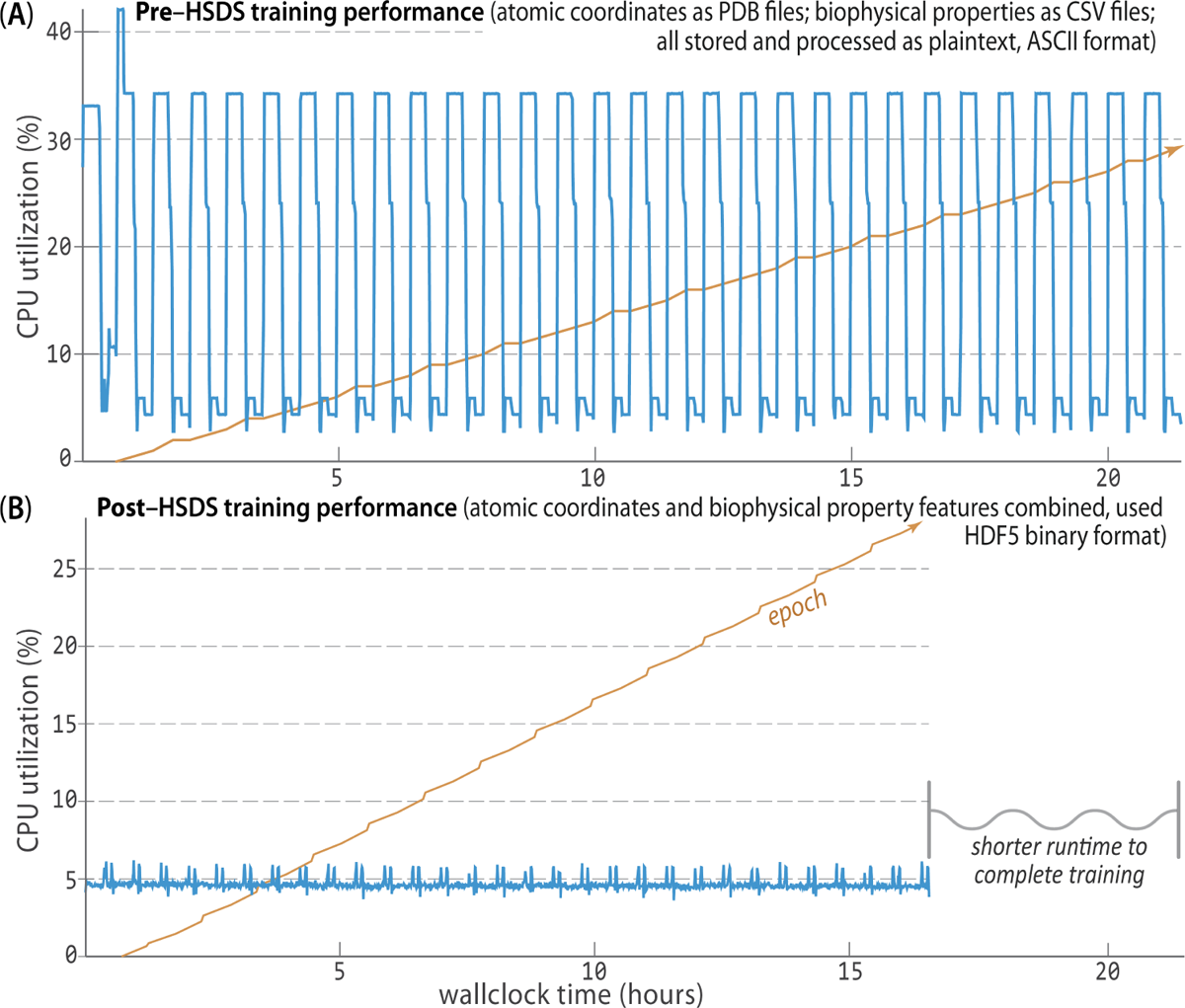
HSDS Affords Significantly Improved Training Runtimes. Using Prop3D, we trained an immunoglobulin-specific variational autoencoder with *≈* 25K domain structures, employing 64 CPUs to process data and four GPUs for 30 epochs of training (orange trace; [12]). **A**. Before we chose to implement HSDS in Prop3D, we stored and processed domain structures as simple plaintext PDB files (parsed with BioPython), along with the corresponding biophysical properties for all atoms in these structures as plaintext files of comma-separated values (CSV; parsed with Pandas). That computation took *≈* 24 hours of wallclock time for *≈* 50K ASCII files on a well-equipped GPU workstation. **B**. Reformulating and streamlining the Prop3D pipeline with HSDS yielded a substantial (*≈* 33%) speed-up: training runtimes across many epochs (orange) improved by *≈* 8 hours (to *≈* 16 hours total), with there being far more efficient CPU usage while reading all of the data (blue traces; note the different vertical scales in **A** and **B**). These data-panel images were exported from our Weights and Biases training dashboard.

### Data Availability & Prop3D’s FAIRness

As summarized in the rest of this section, and detailed in the Supplemental Material (§3), we have sought to make Prop3D FAIR—*Findable*, *Accessible*, *Interoperable*, and *Reproducible* [58]. When possible, the FAIR guidelines would apply both to datasets themselves as well as to the code that underlies the data-generating and data-processing/analysis/reduction pipelines—i.e., a software framework would be FAIR-compliant, insofar as its resultant data are FAIR. Thus, with Prop3D we provide unique identifiers and searchable metadata for open platforms such as Zenodo, WikiData, the Open Science Foundation, and the University of Virginia School of Data Science’s Open Data Portal, as detailed below.

First, the Prop3D-20sf dataset, which contains our prepared structues, pre-computed features and data splits for the 20 highly-populated CATH superfamilies shown in Figure 2, is made available in our HSDS endpoint at the University of Virginia (http://prop3d-hsds.pods.uvarc.io/about) at the domain/CATH/Prop3D-20.h5 (no authentication is necessary; the API must be used as there is not a browser-accessible version). The data can be read into a Python program, as part of one’s ML workflow, using either h5pyd, our Prop3D library. A copy of the raw HDF5 data, exported from our HSDS endpoint, is also available on Zenodo (https://doi.org/10.5281/zenodo.6873024).

The Prop3D library, to run predefined workflows and access our HSDS endpoint, is freely accessible in our GitHub repository (https://github.com/bouralab/Prop3D), with scripts provided to setup HSDS and Kubernetes, e.g. if one plans to run on one’s own local system via k3s.

Finally, all of our Docker-ized tools also can be obtained from our Docker Hub at https://hub.docker.com/u/edraizen.

We have used Wikidata throughout this article to cite the software we use, as well as to create links to the code and data repositories reported herein (e.g., Q108040542 points to Prop3D) [59].

### Summary & Outlook

This work has presented Prop3D, a modular, flexible, Python-based platform that we developed for large-scale protein property featurization and other data-processing/pipelining tasks that typically arise in ML workflows for structural bioinformatics. While Prop3D was developed and deployed as part of a deep learning framework in another project [12], it was intentionally engineered with extensibility and scalability in mind. This tool can be used with local HPC resources as well as in the cloud, and allows one to systematically and reproducibly create comprehensive datasets via the Highly Scalable Data Service (HSDS). Using Prop3D, we have created ‘Prop3D-20sf’ as a new, shared community resource. The Prop3D-20sf protein dataset, freely available as an HSDS endpoint, combines 3D coordinates with biophysical characteristics and evolutionary properties (for each atom), in each structural domain for 20 highly-populated homologous superfamilies in CATH.

The 3D domains in Prop3D-20sf are sanitized via numerous steps, including clean-up of the covalent structure (e.g., adding missing atoms and residues) and physico-chemical properties (protonation and energy minimization). Our database schema mirrors CATH’s hierarchy, mapped to a system based on HDF5 files and including atomic-level features, residue-level features, residue···residue contacts, and pre-calculated train/test/validate splits (in ratios of 80/10/10) for each superfamily derived from CATH’s sequence-identity–based clusters (e.g., ‘S35’ for groups of proteins culled at 35% sequence identity). Notably, our construction of Prop3D-20sf sought to directly and explicitly address the issue of evolutionary data leakage, thereby hopefully mitigating any bias in ML models trained with these datasets. The Prop3D approach and its attendant Prop3D-20sf pre-computed dataset can be used to compare sequence-based (1D), residue-contact–based graphs (2D), and structure-based (3D) methods. For example, one could imagine training a supervised model, with input being a protein sequence, to predict a specific residue-based biophysical property. Similarly, unsupervised models can be trained using one or all of the biophysical properties to learn protein embeddings, such as was the case in our DeepUrfold project [12].

Within Prop3D, we built AtomicToil to enable the facile creation of reproducible workflows, starting with PDB files or by traversing the CATH hierarchy, as well as the Meadowlark toolkit to run Docker-ized structural bioinformatics software. While we primarily developed these tools in order to create the Prop3D-20sf dataset, we envision that the toolkit can be integrated into feature-rich, standalone structural bioinformatics platforms, e.g. BioPython or Biotite. An appealing future direction would be to enable Prop3D’s featurization pipeline to capture information about biomolecular dynamics [60, 61], so as to aid the development of ML models that are more detailed and realistic reflections of protein function. More generally, we believe that Prop3D-20sf and its underlying Prop3D framework may be useful as a community resource in developing workflows that entail processing protein 3D structural information, particularly for projects that arise at the intersection of machine learning and structural bioinformatics.

**Availability and requirements Project name:** Prop3D

**Project home page:** https://github.com/bouralab/Prop3D

**Operating system(s):** Platform independent

**Programming language:** Python

**Other requirements:** Python 3.8 or higher, Singularity or Docker, Toil, Kubernetes

**License:** Creative Commons Attribution 4.0 International License (CC-BY-4).

**Any restrictions to use by non-academics:** None.

## Acknowledgements

We thank Luis Felipe R Murillo (Notre Dame) for technical guidance and help with HSDS at UVA, as well as Lane Rasberry (UVA) for critiquing the manuscript and providing support for Wikidata. We appreciate the early efforts of Menuka Jaiswal, Saad Saleem and Yonghyeon Kweon on this project.

## Abbreviations

| Name | Description/purpose (in this context) | Wikidata entry |
| --- | --- | --- |
| *Structure-based software* |  |  |
| AlphaFold2 | Deep learning-based code for high-accuracy protein 3D structure prediction | Q107711739 |
| AutoDock | A suite of automated protein docking tools | Q4826062 |
| APBS | Adaptive Poisson-Boltzmann Solver, used here to calculate the electrostatic potential for each atom in a given protein | Q65072984 |
| BioPython | General-purpose collection of open-source tools for computational biology | Q4118434 |
| Biotite | A comprehensive library for computational molecular biology | Q114859551 |
| CX | Get curvature for each atom in a given protein | Q114841750 |
| DSSP | Calculate secondary structure and accessibility for each residue in a given structure | Q5206192 |
| EPPIC | Calculate sequence conservation scores for a given protein and obtain biologically relevant protein interactions (i.e., not resulting from crystal packing) | Q114841783 |
| FreeSASA | Get solvent accessibility of each atom in a given protein | Q114841793 |
| MGLTools | Convert atom names to AutoDock names and PDBQT | Q114840701 |
| MODELLER | Create full atom structures from C<sub>α</sub> only models, mutate structures with different amino acids, 'remodel structure' to energy minimize, and model loops | Q3859815 |
| PDB2PQR | Protonate a protein structure, debump hydrogens, energy-minimize, and standardise naming (atomic nomenclature) | Q62856803 |
| pdb-tools | A "Swiss army knife of tools" to manipulate PDB files | Q114840802 |
| SCWRL4 | Correct side-chains using the Dunbrack rotamer library | Q114840881 |
| *Other software* |  |  |
| AWS | Amazon Web Services, on-demand cloud computing platforms | Q456157 |
| Docker | Open-source software for deploying containerized applications | Q15206305 |
| HDF5 | Hierarchical Data Format, version 5 | Q1069215 |
| HSDS | Cloud-native, service-based access to HDF data | Q114859023 |
| h5pyd | Python client library for HDF5 REST interface | Q114859536 |
| Kubernetes | Software to manage containers on a server-cluster | Q22661306 |
| k3s | A light-weight Kubernetes distribution for small servers | Q114860267 |
| MinIO | Cloud storage server compatible with Amazon S3 | Q28956397 |
| NumPy | Numerical programming package for the Python programming language | Q197520 |
| Pandas | Python library for data manipulation and analysis | Q15967387 |
| PyTorch | Open-source, Python-based machine learning library | Q47509047 |
| Toil | Enables creation and deployment of massively parallel workflows in Python | Q114858329 |
| Singularity | Open-source container software for scientific environments | Q51294208 |
| SLURM | Free and open-source job scheduler for Linux and similar (Unix-based) operating systems | Q3459703 |
| Oracle Grid Engine | Supercomputer batch-queuing system | Q2708256 |
| Weights and Biases (wandb) | Python library to track machine learning experiments, version data and manage models | Q107382092 |

### Availability of data and materials

All code is available at https://github.com/bouralab/Prop3D. The ‘Prop3D-20sf’ dataset is available at https://doi.org/10.5281/zenodo.6873024 as a raw HDF5 file, with a public HSDS endpoint at http://prop3d-hsds.pods.uvarc.io/about in domain /CATH/Prop3D-20.h5.

### Ethics approval and consent to participate

Not applicable.

## Consent for publication

Not applicable.

## Authors’ contributions

EJD designed and implemented Prop3D, and drafted/revised the manuscript. JR setup HSDS at UVA and advised on HDF/HSDS best practices. CM advised the work, and drafted/revised the text and figures. PEB advised the overall project. All authors read and approved the final manuscript.

## Author details

^1^Department of Biomedical Engineering, University of Virginia, Charlottesville, VA, USA. ^2^School of Data Science, University of Virginia, Charlottesville, VA, USA. ^3^The HDF Group, Bellevue, WA, USA.

## Supplemental Material for

### 1 Sequence-based bioinformatics tools available in Prop3D

**Table 1:**
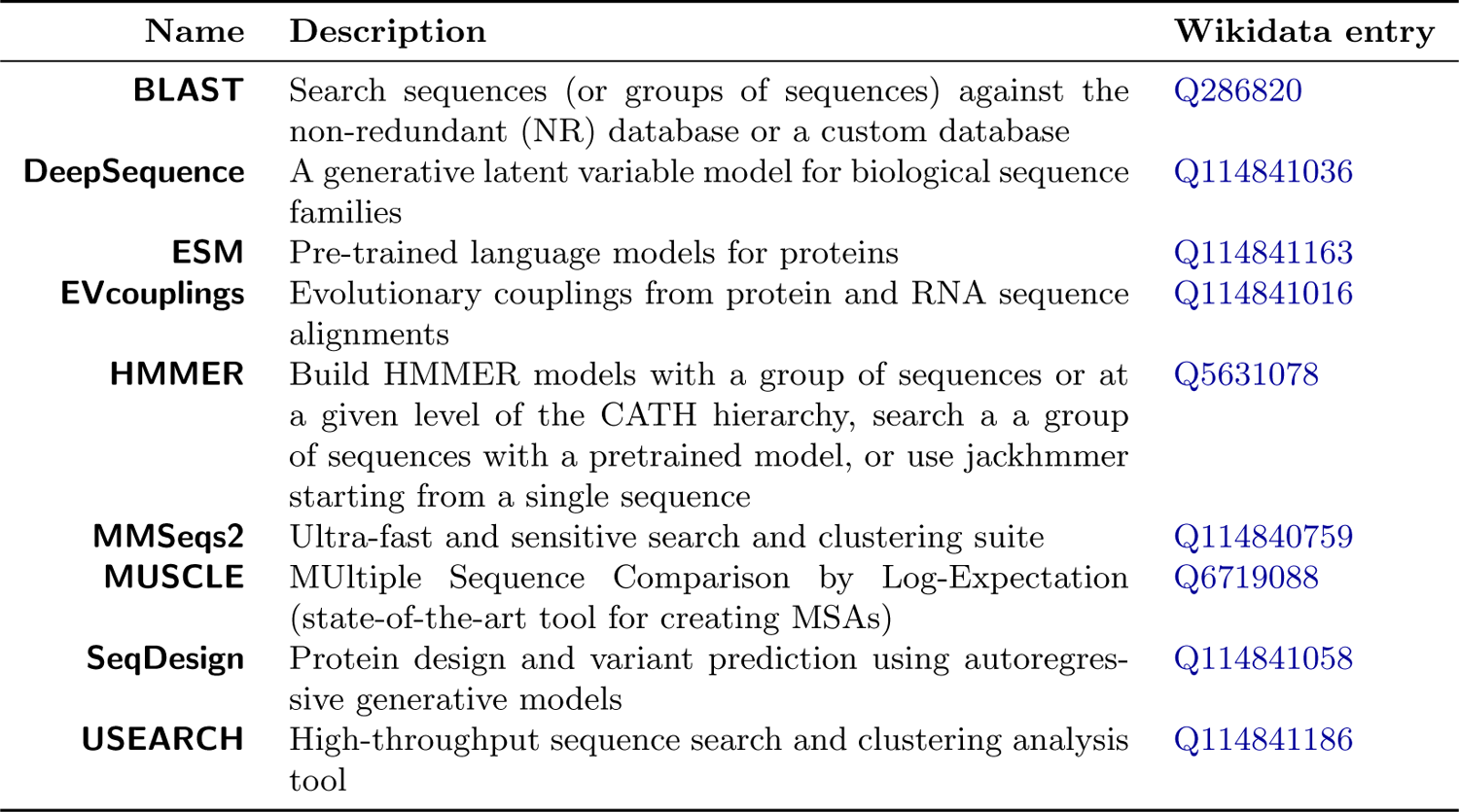
A Catalog of Protein Feature Datasets That Can Be Used in ML. Many different datasets of sequences, structures, and biophysical properties exist. They all contain different amounts of data, data on different levels/scales (chain, domain, residue, atom), and some contain biophysical properties attached to each atom and/or residue. Databases that use atomic coordinates, but without biophysical properties associated with the geometric coordinates, are denoted by daggers (*^†^*).

| Name | Description | Wikidata entry |
| --- | --- | --- |
| **BLAST** | Search sequences (or groups of sequences) against the non-redundant (NR) database or a custom database | Q286820 |
| **DeepSequence** | A generative latent variable model for biological sequence families | Q114841036 |
| **ESM** | Pre-trained language models for proteins | Q114841163 |
| **EVcouplings** | Evolutionary couplings from protein and RNA sequence alignments | Q114841016 |
| **HMMER** | Build HMMER models with a group of sequences or at a given level of the CATH hierarchy, search a a group of sequences with a pretrained model, or use jackhmmer starting from a single sequence | Q5631078 |
| **MMSeqs2** | Ultra-fast and sensitive search and clustering suite | Q114840759 |
| **MUSCLE** | MUltiple Sequence Comparison by Log-Expectation (state-of-the-art tool for creating MSAs) | Q6719088 |
| **SeqDesign** | Protein design and variant prediction using autoregressive generative models | Q114841058 |
| **USEARCH** | High-throughput sequence search and clustering analysis tool | Q114841186 |

### 2 Structural bioinformatics software suites available in Prop3D

**Table 2:** Protein Structure Representations. . Some fundamentally different types of protein structure representations (reps) are schematized here, arranged by dimensionality of the rep. One can always traverse from higher- to lower-dimensional reps without requiring information, while the reverse is not true. Note that some types of reps are more amenable to encapsulation in simple data structures, e.g. protein sequences as character strings (built-in types for programming languages), and residue···residue graphs as adjacency matrices (closely related to *contact maps*). That 3D structures are generally not as ‘cleanly’ representable (in 3D), via available data structures for use in ML workflows, motivates much of Prop3D’s functionality.

| Name | Description/purpose (in this context) | Wikidata entry |
| --- | --- | --- |
| **APBS** | Adaptive Poisson-Boltzmann Solver, used here to calculate the electrostatic potential for each atom in a given protein | Q65072984 |
| **Consurf** | Get pre-calculated conservation scores | Q112888886 |
| **CNS** | Energy minimize a given structure | Q5191443 |
| **CX** | Get curvature for each atom in a given protein | Q114841750 |
| **DSSP** | Calculate secondary structure and accessibility for each residue in a given structure | Q5206192 |
| **EPPIC** | Calculate sequence conservation scores for a given protein and obtain biologically relevant protein interactions (i.e., not resulting from crystal packing) | Q114841783 |
| **foldseek** | Fast searching and clustering of protein structure databases | Q114840749 |
| **FreeSASA** | Get solvent accessibility of each atom in a given protein | Q114841793 |
| **Geometricus** | A structure-based, alignment-free embedding approach for proteins, utilizing moment invariants | Q114840743 |
| **HADDOCK** | Dock two proteins or refine the conformation of two docked proteins | Q114841798 |
| **MaxCluster** | Cluster very similar structures | Q114840623 |
| **MGLTools** | Convert atom names to AutoDock names and PDBQT | Q114840701 |
| **MM-Align** | Align two protein complexes | Q114841843 |
| **mTM-Align** | Multiple structure alignment | Q114841813 |
| **MODELLER** | Create full atom structures from C$_{\alpha}$ only models, mutate structures with different amino acids, ‘remodel structure’ to energy minimize, and model loops | Q3859815 |
| **MSMS** | Calculate molecular surfaces and create meshes | Q114841806 |
| **Multivalue** | Merge electrostatic values from multiple atoms, e.g. on a protein surface | Q114840933 |
| **OpenBabel** | Convert to PDBQT format for AutoDock atom naming | Q612752 |
| **PDB2PQR** | Protonate a protein structure, debump hydrogens, energy-minimize, and standardise naming (atomic nomenclature) | Q62856803 |
| **pdb-tools** | A “Swiss army knife of tools” to manipulate PDB files | Q114840802 |
| **PRODIGY** | Predict binding affinities ($K_D$) and “fraction of common contacts” in complexes | Q114840854 |
| **REDUCE** | Protonate and de-protonate structures | Q114840896 |
| **SCWRL4** | Correct side-chains using the Dunbrack rotamer library | Q114840881 |
| **TM-Align** | Align two or more protein 3D structures | Q114840775 |

### 3 How Prop3D abides by the FAIR guidelines

In general, the creation of scalable, reproducible scientific workflows faces challenges that stem from the sheer volume and heterogeneity of available data-sources and data-types, in addition to potential other factors such as the variable range of computing platforms, architectures and capabilities that one may seek to deploy a workflow across (e.g., multi-core processing on a local workstation, versus a Linux HPC cluster, versus an eScience grid or other highly distributed network environment in the cloud). The ‘FAIR’ principles for scientific data provide a set of best-practices that contribute to the research enterprise by striving to make datasets *Findable*, *Accessible*, *Interoperable*, and *Reproducible*. In other words, FAIR datasets should be (i) easy to *find*, with appropriate metadata to facilitate searching by others; (ii) one should be able to *access* all of the data easily, without undue effort; (iii) one should be able to integrate and otherwise *interoperate* the data with other data-sources and software frameworks; and (iv) the data should be (re)usable and *replicable* by others (a bedrock of the scientific method). When possible, these guidelines (https://www.go-fair.org/fair-principles) would apply equally well to both the datasets themselves as well as to the code that underlies the data-generating and data-processing/analysis/reduction pipelines—i.e., the software framework would be FAIR-compliant, insofar as its resultant data are FAIR. The following enumerates how Prop3D complies with these guidelines:

#### 1. Findable

The first step in (re)using data is to be able to find it. Metadata and data should be easy to find for both humans and computers. Machine-readable metadata are essential for automated discovery of datasets and services, so this is an essential component of the FAIRification process. In Prop3D, the following hold:

F1. (Meta)data are assigned a globally unique and persistent identifier

F2. Data are described with rich metadata (defined by R1 below)

F3. Metadata clearly and explicitly include the identifier of the data they describe

F4. (Meta)data are registered or indexed in a searchable resource

#### 2. Accessible

Once a user finds the required data, she/he/they need to know how such data can be accessed, possibly including issues of authentication and authorisation. In Prop3D,

A1. (Meta)data are retrievable by their identifier using a standardised communications protocol

A1.1 The protocol is open, free, and universally implementable

A1.2 The protocol allows for an authentication and authorisation procedure, where necessary

A2. Metadata are accessible, even when the underlying data are no longer available

#### 3. Interoperable

Datasets generally are not used in a vacuum, and at some point or another will need to be integrated with other types and sources of data. In addition, the data need to interoperate with available applications or workflows for analysis, storage, and processing. In Prop3D,

I1. (Meta)data use a formal, accessible, shared, and broadly applicable language for knowledge representation.

I2. (Meta)data use vocabularies that follow FAIR principles

I3. (Meta)data include qualified references to other (meta)data

#### 4. Reusable

The ultimate goal of FAIR is to optimise the reuse of data. To achieve this, metadata and data should be well-described so that they can be replicated and/or combined in different settings. In Prop3D,

R1. (Meta)data are richly described with a plurality of accurate and relevant attributes

R1.1. (Meta)data are released with a clear and accessible data usage license

R1.2. (Meta)data are associated with detailed provenance

R1.3. (Meta)data meet domain-relevant community standards

